# The Exon Junction Complex Core Represses Caner-specific Mature mRNA Re-splicing: A Potential Key Role in Terminating Splicing

**DOI:** 10.1101/2021.04.01.438154

**Authors:** Yuta Otani, Ken-ichi Fujita, Toshiki Kameyama, Akila Mayeda

## Abstract

Using the TSG101 pre-mRNA, we previously discovered cancer-specific resplicing of mature mRNA that generates aberrant transcripts/proteins. The fact that mRNA is aberrantly re-spliced in various cancer cells implies there must be an important mechanism to prevent deleterious re-splicing on the spliced mRNA in normal cells. We thus postulated that the mRNA re-splicing is controlled by specific repressors, and we searched for repressor candidates by siRNA-based screening for mRNA re-splicing activity. We found that knockdown of EIF4A3, which is a core component of the exon junction complex (EJC), significantly promoted mRNA re-splicing. Remarkably, we could recapitulate cancer-specific mRNA resplicing in normal cells by knock-down of any of the core EJC proteins, EIF4A3, MAGOH or RBM8A (Y14), implicating the EJC core as the repressor of mRNA re-splicing often observed in cancer cells. We propose that the EJC core is a critical mRNA quality control factor to prevent over-splicing of mature mRNA.

## 1. INTRODUCTION

Although the basic mechanism of how pre-mRNA splicing is initiated and proceeded has been characterized in detail, it is less well known about how splicing is terminated. *Cis*-acting splicing signals, such as 5’/3’ splice sites and branch site followed by poly-pyrimidine tract, are not highly conserved, and thus they are necessary but not sufficient to be utilized [reviewed in 1]. Because of this redundancy, there remain many splice site-like sequences in fully spliced mRNA. However, these spurious splice sites are usually never used, and the mature mRNA is safely exported to the cytoplasm for translation. These processes are definitely critical to maintain the integrity of coding mRNA. Whereas in the vertebrate multi-step splicing pathways, such as recursive splicing [2,3], intrasplicing [4], and nested splicing [5,6], it is evident that the initially spliced intermediates can serve to be spliced again, indicating that the splicing process is not terminated when certain conditions are met. One component in the termination process could be the DEAH-box family RNA helicases involved in the dissociation of spliceosome, such as DHX8 (also known as HRH1, PRP22) and DHX15 (PRP43) [reviewed in 7]. However, the exact mechanism to control splicing termination is poorly understood. We address this thorny problem by taking advantage of an aberrant multistep splicing. In this event, a normally spliced mRNA can also be a substrate for further splicing often in cancer cells, which is termed cancer-specific mRNA re-splicing.

Aberrant splicing that occurs over very long distances (skipping many exons with flanking huge introns) to generate annotated truncated mRNAs, are often observed in cancer cells [reviewed in 8]. TSG101 and FHIT pre-mRNAs, for instance, were often reported as typical cases. Using these two substrates, we discovered that the mRNA re-splicing event occurs on mature spliced mRNA, in which the normal splicing removes all authentic splice sites and bringing the weak re-splice sites into close proximity [9]. The re-spliced TSG101 mRNA, TSG101Δ154-1054 mRNA (Figure 1a), produces an aberrant protein product that up-regulates the full-size TSG101 by interfering with the ubiquitin-dependent degradation of TSG101 protein [10]. Importantly, the overexpression of TSG101 protein enhances cell proliferation and promotes malignant tumor formation in nude mice [10,11]. TSG101 is essential functions for cell proliferation and survival, while its overexpression is closely associated with aggressive metastasis and progression in various cancers [Reviewed in 12]. It is thus critical to control this kind of aberrant mRNA re-splicing pathway in normal cells to prevent production of potentially deleterious protein products.

**Figure 1.**
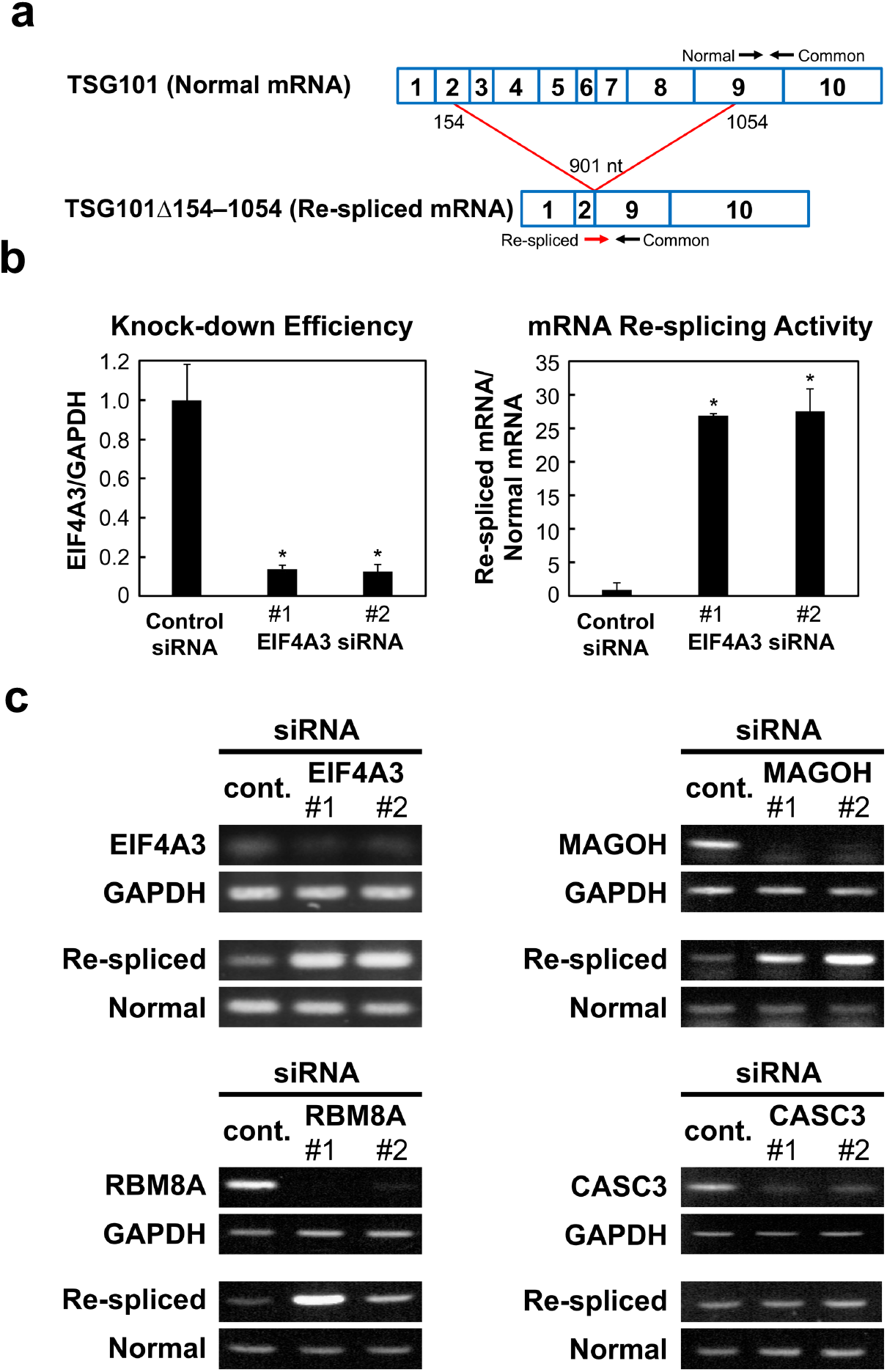
Identification of EJC-core factors as repressor candidates of cancer-specific mRNA re-splicing. (**a**) The schematic structure of normally spliced TSG101 mRNA and cancer-specifically re-spliced TSG101Δ154–1054 mRNA. The primer sets for the detection were indicated with arrows. (**b**) To measure the generated normal TSG101 and re-spliced TSG101Δ154-1054 mRNAs, the extracted total RNAs were analyzed by RT–qPCR using the indicated specific primer sets in (A). To knock-down the expression of the endogenous *EIF4A3* gene, MCF-7 cells were transfected with EIF4A3 siRNAs together with control siRNA. The cell extracts were then analyzed by RT–qPCR and the relative amounts of EIF4A3 mRNA are plotted. The histograms represent the means ± standard deviations of three replicates (\**P* < 0.05). (**c**) To knock-down the expression of endogenous EJC core factor genes, MCF-7 cells were transfected with siRNA targeting EIF4A3, MAGOH, RBM8A, CASC3 or control siRNA (cont.). To observe the knocked-down mRNAs and generated normal TSG101 and re-spliced TSG101Δ154-1054 mRNAs, the extracted total RNAs were analyzed by RT– PCR The depletion of the targeted proteins was verified by immunoblotting (Figure S2).

Here we postulated that the control of the aberrant mRNA re-splicing is promoted by a specific repressor in normal cells. To test this hypothesis, we performed siRNA screening of repressor candidates based on cancer-specific mRNA re-splicing activity of the TSG101 pre-mRNA. This approach identified EIF4A3, whose knock-down significantly stimulated re-splicing of the TSG101 mRNA. EIF4A3 is one of three essential core components of the exon junction complex (EJC) along with MAGOH and RBM8A (Y14). The EJC is deposited upstream of the spliced exon-exon junctions as a consequence of pre-mRNA splicing; and the EJC together with the recruited peripheral factors play important roles in multiple post-splicing events, such as mRNA export, nonsense-mediated mRNA decay (NMD), and translation [reviewed in 13,14]. Here we demonstrate that EJC core *per se*, but not the EJC peripheral factors, is necessary to repress mRNA re-splicing in normal cells.

## 2. RESULTS

### 2.1. Re-splicing of mRNA Is Enhanced by EIF4A3 Knock-down

Previously, it was demonstrated that the induction and overexpression of the tumor suppressor gene *TP53* (*p53*) prevents cancer-specific aberrant splicing of TSG101 pre-mRNA [10,15], i.e., generation of the re-spliced TSG101Δ154-1054 mRNA [9]. Therefore, this mRNA re-splicing event is under the control of TP53, presumably indirectly through other regulators; i.e., upregulation of an mRNA re-splicing repressor and/or down-regulation of an mRNA re-splicing activator. To identify such repressor or activator candidates, we activated TP53, using highly selective MDM2 inhibitor RG7388 [16] to repress aberrant re-splicing and searched for up- or down-regulated genes on a microarray (Tables S1). We confirmed that the activation of TP53 by RG7388 indeed prevents re-splicing activity of TSG101 pre-mRNA (Figure S1).

We selected 48 candidate proteins including hnRNP proteins, RNA helicases, a splicing-related kinase, and other RNA-binding proteins (Table S2) from the up- and down-regulated genes upon activation of TP53. Then we performed siRNA-mediated knock-down of each protein (Table S2 for the siRNA sequences) in the human metastatic mammary carcinoma cell line MCF-7, in which mRNA re-splicing was previously demonstrated [9]. We found that the knocking down EIF4A3 markedly increased the aberrant re-spliced product of TSG101 mRNA (TSG101Δ154-1054) over twenty times more than in the control (Figure 1b). Our results indicate that EIF4A3 is a potent repressor of mRNA re-splicing. Since EIF4A3 expression was not up-regulated, but rather down-regulated, under the RG7388-mediated activation of TP53 (Table S1), EIF4A3 was evidently not a TP53-dependent repressor of mRNA re-splicing (see Discussion).

### 2.2. The EJC Core Factors Were Identified as Repressors of mRNA Re-splicing

Remarkably, EIF4A3 protein, which appers to repress mRNA re-splicing, is one of the core components of the EJC that is deposited onto mature mRNA [reviewed in 13,14]. Next, we test whether EIF4A3 functions solely or EIF4A3 functions as a core component in the EJC. We thus knocked-down the other EJC core components, MAGOH, RBM8A and CASC3 (MLN51) using siRNAs in MCF-7 cells.

Like for EIF4A3, we found that knock-down of both MAGOH and RBM8A markedly activated the aberrant re-splicing of TSG101 mRNA, whereas CASC3 knock-down had no effect (Figure 1c). Since EIF4A3, MAGOH and RBM8A are essential, but CASC3 is dispensable, for the assembly of a stable EJC core [17], it is reasonable to assume that an intact EJC core assembly is necessary to repress mRNA re-splicing. We confirmed the actual binding of the EJC core on TSG101 mRNA by RNA-immunoprecipitation assay using anti-RBM8A antibody (Figure S3). Together, we conclude that the formation of the EJC core per se is a substantial repressor of mRNA re-splicing.

### 2.3. The EJC Core Represses mRNA Re-splicing in Normal Cells

If the EJC core alone is necessary to repress mRNA re-splicing in normal cells, the knock-down of each EJC core factor in normal cells should activate mRNA re-splicing as observed in cancer cells.

We tested this assumption using MCF-10A cells, which is a non-cancerous human mammary epithelial cell line. We could not detect re-spliced mRNA product in MCF-10A cells (Figure 2a), just as we had observed in normal mammary epithelial cells [9]. Remarkably, mRNA re-spliced product was generated under efficient siRNA-mediated knock-down of each EJC core factor, EIF4A3, MAGOH or RBM8A (Figure 2b). We conclude that knockdown of a single EJC core factor is sufficient to activate mRNA re-splicing in normal cells.

**Figure 2.**
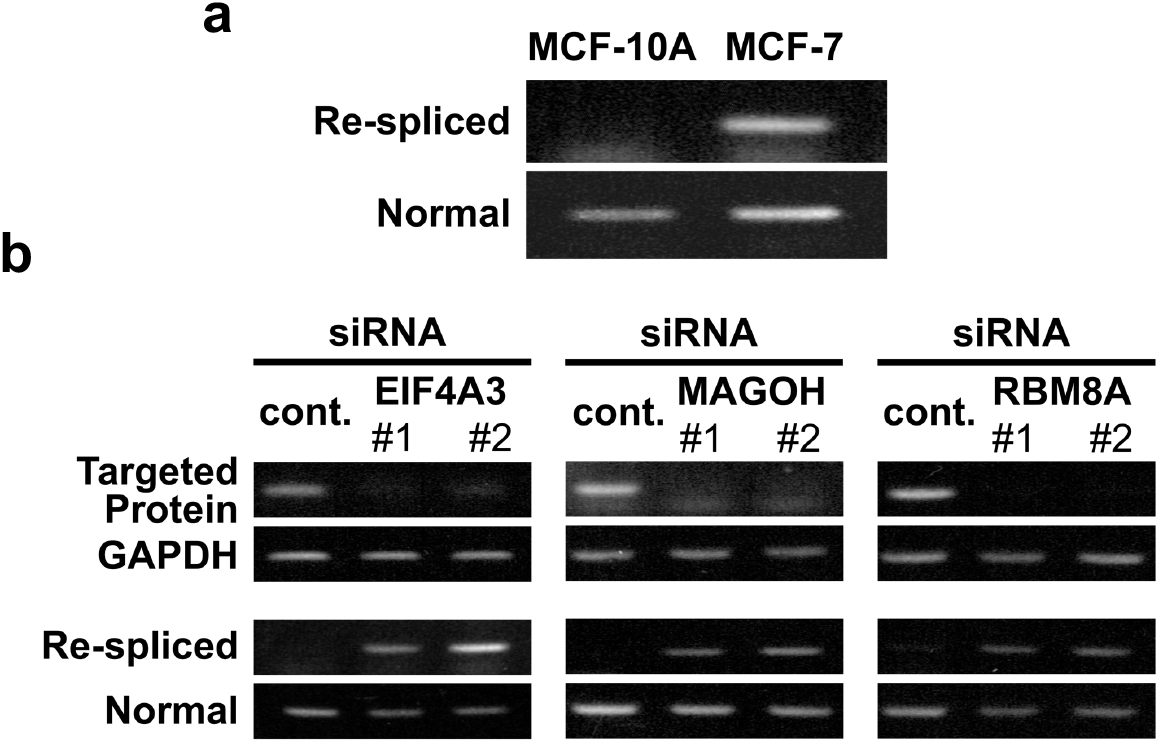
Depletion of EJC-core factors can induce cancer-specific mRNA re-splicing in non-cancerous MCF-10A cells. (**A**) Total RNAs extracted from MCF-10A or MCF-7 cells were analyzed by RT-PCR using specific primer sets (see Figure 1b). (**B**) To knock-down the expression of endogenous EJC core factor genes, MCF-10A cells were transfected with the indicated siRNAs. To observe the knocked-down efficiency and TSG101 re-splicing activity, the extracted total RNAs were analyzed by RT–PCR. The depletion of the targeted proteins was verified by immunoblotting (Figure S2).

### 2.4. mRNA Re-splicing Is Not Due To Repression of EJC-mediated NMD

The EJC contains peripheral factors responsible for nonsense-mediated mRNA decay (NMD) such as the UPF (UP-Frameshift) proteins (UPF1, 2, 3) [reviewed in 13,14]. As the re-spliced product, TSG101Δ154-1054 mRNA, contains a premature termination codon (PTC) downstream of re-spliced junction, it could be a target of NMD [9]. Therefore, knock-down of EJC core factors could also dissociate UPF proteins so that it potentially represses NMD of re-spliced mRNA leading to its eventual increase.

Mature mRNA re-splicing, generating TSG101Δ154-1054 mRNA, is a cancer-specific event and it is barely detectable in normal cells [9; and references therein]. First, it is critical to determine in normal cells whether the re-spliced mRNA is indeed not generated or it is the result of the degradation of the generated mRNA by NMD. We thus performed siRNA-mediated knock-down of essential NMD factor UPF1 in normal MCF-10A cells. This knock-down effectively depleted UPF1 but it did not generate the re-spliced TSG101Δ154-1054 product (Figure 3a), confirming that the re-spliced TSG101Δ154-1054 product barely exists in normal cells, and indicating that this is not due to NMD.

**Figure 3.**
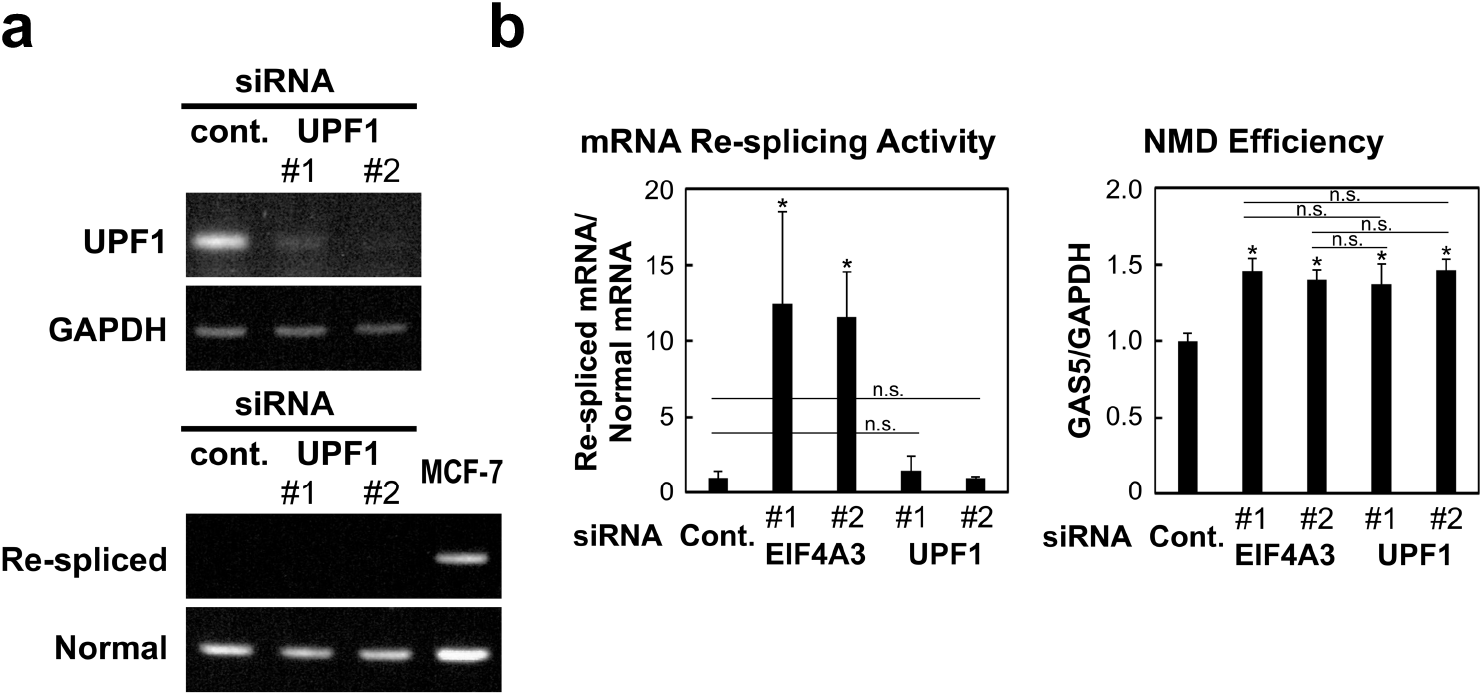
Activation of mRNA re-splicing by the knock-down of EJC core factors is not due to the repression of EJC-mediated NMD. (**a**) To knock-down the expression of endogenous NMD factor UPF1, MCF-10A cells were transfected with the indicated siRNAs. To observe the knocked-down mRNA and TSG101 re-splicing activity, the extracted total RNAs were analyzed by RT–PCR. MCF-7 cells (control) shows the re-spliced mRNA that lacks in other lanes using MCF-10A cells. (**b**) HeLa cells were transfected with EIF4A3 or UPF1 siRNAs (or control siRNA). To evaluate the re-splicing activity with TSG101 mRNA and the NMD efficiency with GAS5 mRNA, the extracted total RNAs were analyzed by RT– qPCR using specific primer sets. The histograms represent the means ± standard deviations of three replicates (\**P* < 0.05, n.s.=not statistically significant *P* > 0.05).

We therefore quantified mRNA re-splicing activity under the knock-down of UPF1 and EIF4A3 using HeLa cells (in which siRNA-mediated interference is more efficient than that in MCF-10A cells). The mRNA re-spliced product was markedly increased by the depletion of EIF4A3, but not at all by the depletion of UPF1 (Figure 3b, left graph). In this assay, the level of GAS5 mRNA, a well-known endogenous NMD target [18,19], was significantly increased in both EIF4A3 and UPF1 depleted cells (Figure 3b, right graph), supporting that NMD is inhibited almost equally by knocking down of either EIF4A3 or UPF1. Together, we conclude that the observed increase of mRNA re-spliced product by the knock-down of EJC core factor is attributed to the intrinsic function of EJC as a repressor of mRNA re-splicing, but is not due to the repression of EJC-mediated NMD.

### 2.5. mRNA Re-splicing Is Not Due To Repression of EJC-mediated mRNA Export

As well as several NMD factors, the EJC also interacts peripherally with several mRNA export factors and splicing regulators [reviewed in 13,14]. It is conceivable that knock-down of EJC core factor could dissociate mRNA export factor such as NXF1 (TAP), which might lead to the nuclear retention of mature mRNA. Then the nuclear-retained mRNA could be a target of post-transcriptional re-splicing. It is also possible that the dissociation of other EJC-associated peripheral factors such as ACIN1 (ACINUS), PNN (Pinin), RNPS1, and SAP18, known as potential splicing regulators, could affect the efficiency of mRNA re-splicing.

We thus tested these EJC peripheral factors by siRNA-mediated knock-down in MCF-7 cells (Figure 4a). The mRNA re-splicing was not significantly activated (Figure 4b) as we observed in the depletion of EJC core factors (Figure 1b, c). The knock-down of other EJC-associated mRNA export factors, UAP56 and URH49, also had no significant effects on mRNA re-splicing activity (unpublished results). Our results clearly implicate that mRNA re-splicing event is not due to the over-splicing on the nuclear retained mRNA caused by inhibition of mRNA export. We conclude that the formed EJC core itself, but not the associated EJC-peripheral factors, is involved in the prevention of mRNA re-splicing.

**Figure 4.**
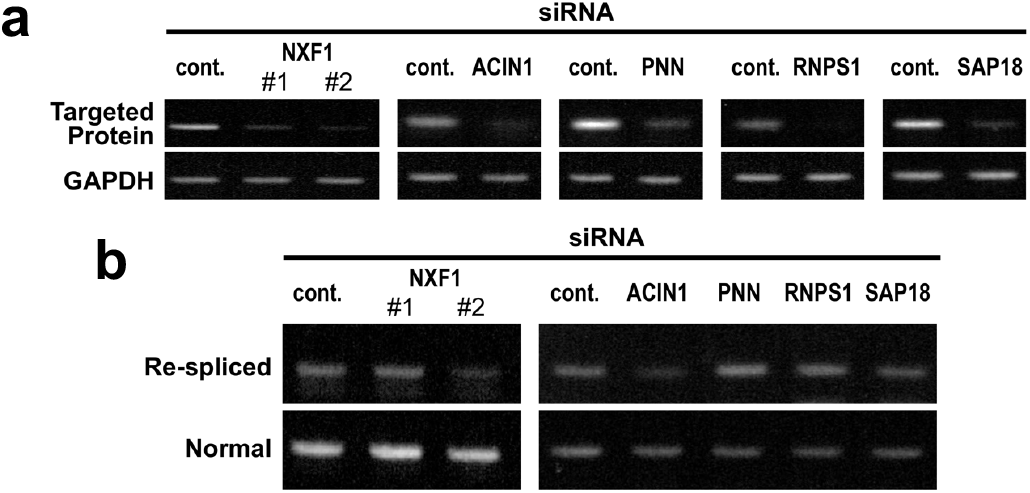
Depletion of EJC-peripheral factors, including mRNA export factor, have no significant effects on mRNA resplicing in MCF-7 cells. (**a, b**) To knock-down the expression of endogenous EJC-peripheral factors (NXF1, ACIN1, PNN, RNPS1, SAP18), MCF-7 cells were transfected with the indicated siRNAs. To observe the knocked-down efficiency and TSG101 re-splicing activity, the extracted total RNAs were analyzed by RT–PCR.

### 2.6. RBM8A Expression is Relevant to Cancer-specific mRNA Re-splicing Activity

TSG101 mRNA is frequently re-spliced in various cancer tissues for unknown reasons. Since we have identified the EJC core factor as a repressor of mRNA re-splicing, it is reasonable to predict reduced expression of an EJC core factor in cancer cells. We indeed observed a modest decrease of RBM8A protein in cancer MCF-7 cells compared to that in non-cancerous MCF-10A cells (Figure 5a, b). Remarkably, overexpression of RBM8A in stable cell line significantly restored the repression of mRNA re-splicing (Figure 5c, d), indicating that the observed repression in MCF-7 cells is partly due to down-regulation of RBM8A.

**Figure 5.**
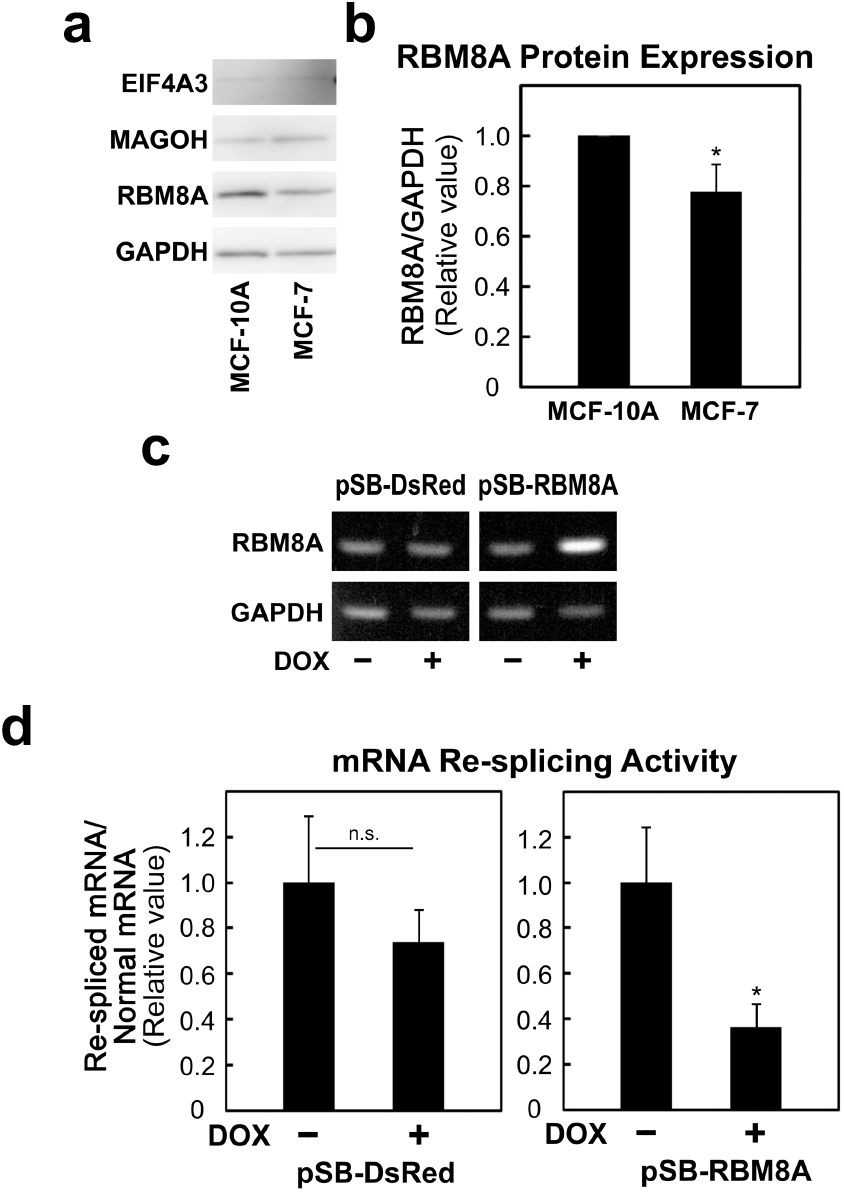
RBM8A protein expression is lower in cancer cells as compared with that in non-cancerous cells, and RBM8A overexpression represses mRNA re-splicing in cancer cells. (**a**) Whole cell extracts from MCF-10A and MCF-7 cells were analyzed by immunoblotting with the indicated specific antibody. GAPDH was used as an endogenous control. (**b**) RBM8A protein expression was normalized to that of GAPDH and quantified by densitometry. The histograms represent the means ± standard deviations of three replicates (\**P* < 0.05). (**c**) The stable MCF-7 cell lines expressing control and RBM8A proteins (pSB-DsRed and pSB-RBM8A) were cultured with or without doxycycline (DOX) for 48 h. The total RNAs were analyzed by RT–PCR using the indicated primer sets. (**d**) To evaluate the re-splicing activity with TSG101 mRNA, the total RNAs were analyzed by RT-qPCR using specific primer sets. The histograms represent the means ± standard deviations of three replicates (\**P* < 0.05, n.s. *P* > 0.05).

## 3. DISCUSSION

Here we demonstrate that the minimum three core EJC factors, without any EJC peripheral factors, is responsible for repressing deleterious re-splicing on mature mRNA in normal cells, implicating the novel role of EJC core on mature mRNA in terminating constitutive splicing (Figure 6). On the other hand, two intriguing questions remain; namely, why mRNA re-splicing often occurs in cancer cells but not in normal cells, and what is the exact mechanism underlying the EJC-triggered repression of mRNA re-splicing; which are the main limitations of this study are that further work is necessary to answer the open questions, but they are worthwhile discussing in this communication but are worthwhile discussing in this communication.

**Figure 6.**
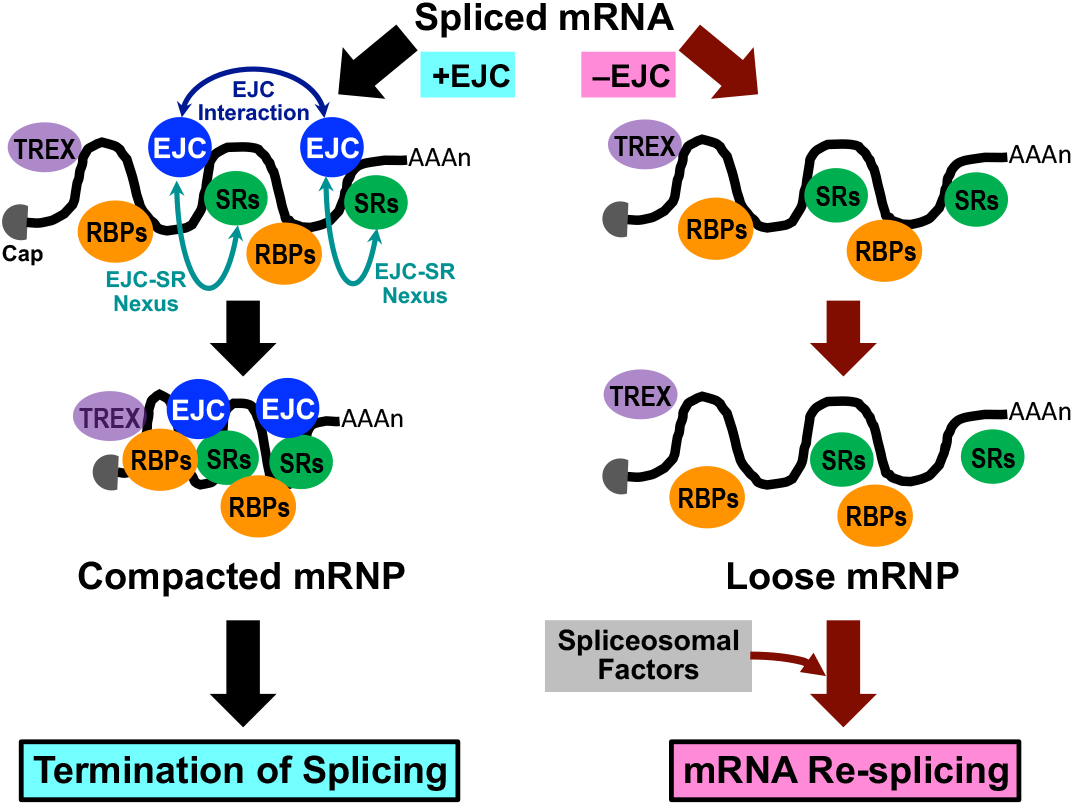
Model of the EJC-triggered pathway leading to the termination of splicing. The spliced mRNA was shown to form packed and compacted structure of mRNP by bindings between EJC cores and among EJC and SR proteins (EJC-SR nexus) together with other RNA-binding proteins (RBPs) [20,21]. We assume that splicing-dependent binding of EJC core to mature mRNA is essential to initiate the formation of this highly compacted mRNP structure.

Using MCF-7 breast cancer cells as a model, we showed that the down-regulation of EJC core factor, RBM8A, is one of the causes of mRNA re-splicing. We need extensive investigations to examine whether the repression of mRNA re-splicing is due to a scarcity of viable EJC cores using different kinds of cancer cells. Since EIF4A3 was not a TP53-dependent re-splicing repressor, there could be another unidentified regulator of mRNA re-splicing, which is under the control of the tumor suppressor gene *TP53*. All these repressors, together with conceivable cancer-specific resplicing activators should be comprehensively investigated to elucidate the cause of mRNA resplicing frequently observed in various cancer cells/tissues (Figure S4).

Regarding the involved mechanism in the EJC-triggered repression of mRNA re-splicing, we assume the following two scenarios; Hypothesis 1: The non-canonical binding of the ECJ core to the corresponding alternative splice sites on the spliced mRNA causes steric hindrance to prevent mRNA re-splicing. Hypothesis 2: The EJC promotes spliced mRNA into highly compacted structure of mRNA particles (mRNPs) with other proteins that interfere with re-assembly of early spliceosome to initiate mRNA re-splicing (Figure 6).

Considering the ‘Hypothesis 1’, recent studies uncovered EJC-triggered PSAP (complex with PNN, SAP18 and RNPS1)-dependent and PSAP-independent mechanisms that repress the use of spurious and recursive splice sites (Figure S5) [22,23]. According to this PSAP-independent mechanism, which is our case in the repression of mRNA re-splicing (Figure 4), the EJC core must directly bind upstream of the 3’ re-splice sites to prevent re-splicing [22]. However, reported data of cross-linking and immunoprecipitation coupled to high-throughput sequencing (CLIP-Seq) in TSG101 mRNA indicated that the significant binding sites of the EJC core factor, EIF4A3, were located more than 18 nucleotides downstream from the 3’ re-splice sites [24]. Therefore, the possibility of masking the re-spliced 3’ splice sites by the EJC core is unlikely. (Figure S5).

The ‘Hypothesis 2’ is more promising. After the release of the spliced mRNA from the spliceosome by the helicase DHX8, bare mature mRNA has a risk of nuclease digestion. Biochemical analysis of the components of mature mRNPs revealed that EJCs multimerize together with other RNA-binding proteins including SR proteins to form mega-Dalton sized higher-order complexes that protect spliced mRNA from endonuclease digestion [20,21]. This fact prompted us to assume that the EJC core depletion could prevent the formation of such highly compacted structures on spliced mRNA so that it allows re-assembly of the dissociated spliceosome to initiate mRNA re-splicing (Figure 6). Further work is under way to address this attractive hypothesis.

It is conceivable that the EJC core is a common key factor controlling different types of the multistep splicing process [22,23]. Using TSG101 pre-mRNA as a model substrate of cancer-specific aberrant mRNA re-splicing, we demonstrate that the deposition of the EJC safeguards against the deleterious over-splicing event. We propose that EJC functions as a key signal to terminate pre-mRNA splicing and thus plays a further role to maintain quality of the protein-coding transcriptome (Figure S4).

## 4. MATERIALS AND METHODS

### 4.1. Cell Culture and siRNA-mediated Knock-down

MCF-7 cells, obtained from the Cell Resource Center (http://www2.idac.tohoku.ac.jp/dep/ccr/index_en.html), were cultured in Dulbecco’s modified Eagle’s medium (Fujifilm Wako Chemicals, Kyoto, Japan) supplemented with 10% fetal bovine serum (Sigma-Aldrich, St. Louis, MO, USA). MCF-10A cells, obtained from the American Type Culture Collection (ATCC), were cultured in a special medium (MEGM with MEGM SingleQuots; Lonza, Walkersville, MD, USA).

MCF-7 and MCF-10A cells were transiently transfected with the indicated siRNAs (Table S3) using Lipofectamine RNAi MAX (Thermo Fisher Scientific, Waltham, MA, USA), according to the manufacturer’s instructions. At 48 h post-transfection, total RNA was isolated from siRNA-treated cells using an RNeasy Mini Kit (QIAGEN, Venlo, Nederland) according to the manufacturer’s instructions. Degradation of the targeted mRNAs was analyzed by RT–PCR (see Section 4.3) and depletion of the protein products was verified by immunoblotting (see Section 4.4).

### 4.2. Establishment of Stable MCF-7 Cell Lines and Induction of RBM8A

All the synthetic oligonucleotides used as primers were purchased (Integrated DNA Technologies, Tokyo, Japan). The coding-fragment of RBM8A protein was amplified by PCR on cDNA prepared from MCF-7 cells using KOD-Plus-Neo polymerase (Toyobo, Osaka, Japan) with the specific primer set (Table S3). The amplified DNA was subcloned into a TA cloning vector pGEM-T (Promega, Madison, WI) using TaKaRa Ex Taq polymerase (Takara Bio, Kusatsu, Japan). The obtained pGEM-T-RBM8A plasmid was cleaved with Sfi I and subcloned into the Sfi I site of pSBtet-GP vector (Addgene, Watertown, MA, USA) [25].

In order to establish stable cell lines, the pSBtet-GP constructs (pSB-DsRed and pSB-RBM8A) were co-transfected with the Sleeping Beauty transposon vector pCMV(CAT)T7-SB100X (Addgene) [26] using Lipofectamine 3000 (Thermo Fisher Scientific), according to the manufacturer’s instructions. At 48 h post-transfection, cells were selected with 1 μg/ml puromycin (Sigma-Aldrich) for ~10 days and terminated when most of cells showed GFP-derived green fluorescence. To induce RBM8A and the control DsRed proteins, 50 ng/ml of doxycycline (Takara Bio USA, Mountain View, CA, USA) were added to each cell lines and cultured for 48 h.

### 4.3. RT–PCR and RT–qPCR Analyses

The cDNA was synthesized from total RNA using SuperScript III reverse transcriptase (Thermo Fisher Scientific) with Oligo(dT)18 primer (Thermo Fisher Scientific), and analyzed by PCR using TaKaRa Ex Taq polymerase (Takara Bio) with target-specific primers (Integrated DNA Technologies, Coralville, IA, USA; Table S3). The PCR products were visualized by 2% agarose gel electrophoresis. The PCR products were further quantified by Applied Biosystems 7900HT Fast Real Time PCR System (Thermo Fisher Scientific) using THUNDERBIRD SYBR qPCR Mix (Toyobo).

### 4.4. Immunoblotting Analysis

Cells were lysed with RIPA buffer supplemented with Protease Inhibitor Cocktail (Nacalai Tesque, Kyoto, Japan) according to the manufacturer’s instructions. Then, the cell lysates were separated by 15% SDS-polyacrylamide gel electrophoresis and electro-blotted onto a Immun-Blot PVDF Membrane using a Trans-Blot SD Semi-Dry Transfer Cell (Bio-Rad, Hercules, CA, USA). The membranes were blocked with 2% skimmed milk and probed with antibodies, essentially as described [27]. The primary antibodies against GAPDH (1:2000 dilution; Medical & Biological Laboratories, Nagoya, Japan), EIF4A3 (1:1000; Abcam, Cambridge, UK), RBM8A (1:1000; GeneTex, Irvine, CA, USA), and MAGOH (1:1000; Medical & Biological Laboratories, Tokyo, Japan) were used. The secondary antibodies were the anti-rabbit IgG-HRP (1:2500; Sigma-Aldrich) and anti-mouse IgG-HRP (1:2500; Sigma-Aldrich). Immuno-reactive proteins were visualized using the Immobilon Western Chemiluminescent HRP Substrate (Merck Millipore, Burlington, MA, USA). The detected bands on the scanned blots were quantitated with ImageJ freeware.

### 4.5 RNA Immunoprecipitation (RIP) Assay

The RIP assay with MCF-7 cells was performed using the RIP-Assay Kit (Medical & Biological Laboratories) with anti-RBM8A antibody (Medical & Biological Laboratories) according to the manufacturer’s instructions. The immunoprecipitated RNAs were amplified by RT–PCR using the primers targeting TSG101 mRNA (Table S3) and visualized by 2% agarose gel electrophoresis.

### 4.6. Statistical Analysis

Analyzed expression levels from two groups were compared using Welch’s t-test. One-way analysis of variance (ANOVA) followed by post-hoc tests were used to analyze results among multiple groups with a normal distribution and equal variance. Statistical significance was set at P < 0.05. Data were presented as mean ± standard deviations.

## DATA AVAILABILITY STATEMENT

The original microarray data (see Table S1) have been deposited in the NCBI’s Gene Expression Omnibus (GEO) database under accession number GSE178102.

## AUTHOR CONTRIBUTIONS

Organizer of project, A.M.; plan of research, Y.O., T.K., and A.M.; most of experiments, Y.O.; additional experiments for Supplementary figure, K.-i.F.; data analyses, Y.O., T.K., K.-i.F., and A.M.; writing—original draft preparation, Y.O.; writing—review and editing, Y.O., T.K., K.-i.F., and A.M.; project supervision and administration, A.M. All authors have read and agreed to the published version of the manuscript.

## ACKNOWLEDGMENTS

We are grateful to H. Naito, K. Takagaki, H. Fujiwara, and A. Matsuura in Nippon Shinyaku Co., Ltd. for their support and encouragements for Y.O. We thank all members of the Mayeda and Ishi laboratories for their constructive discussions, and Drs. N. Kataoka and J. Venables for critical reading of the manuscript.

## Supplementary Materials

**Table S3.**
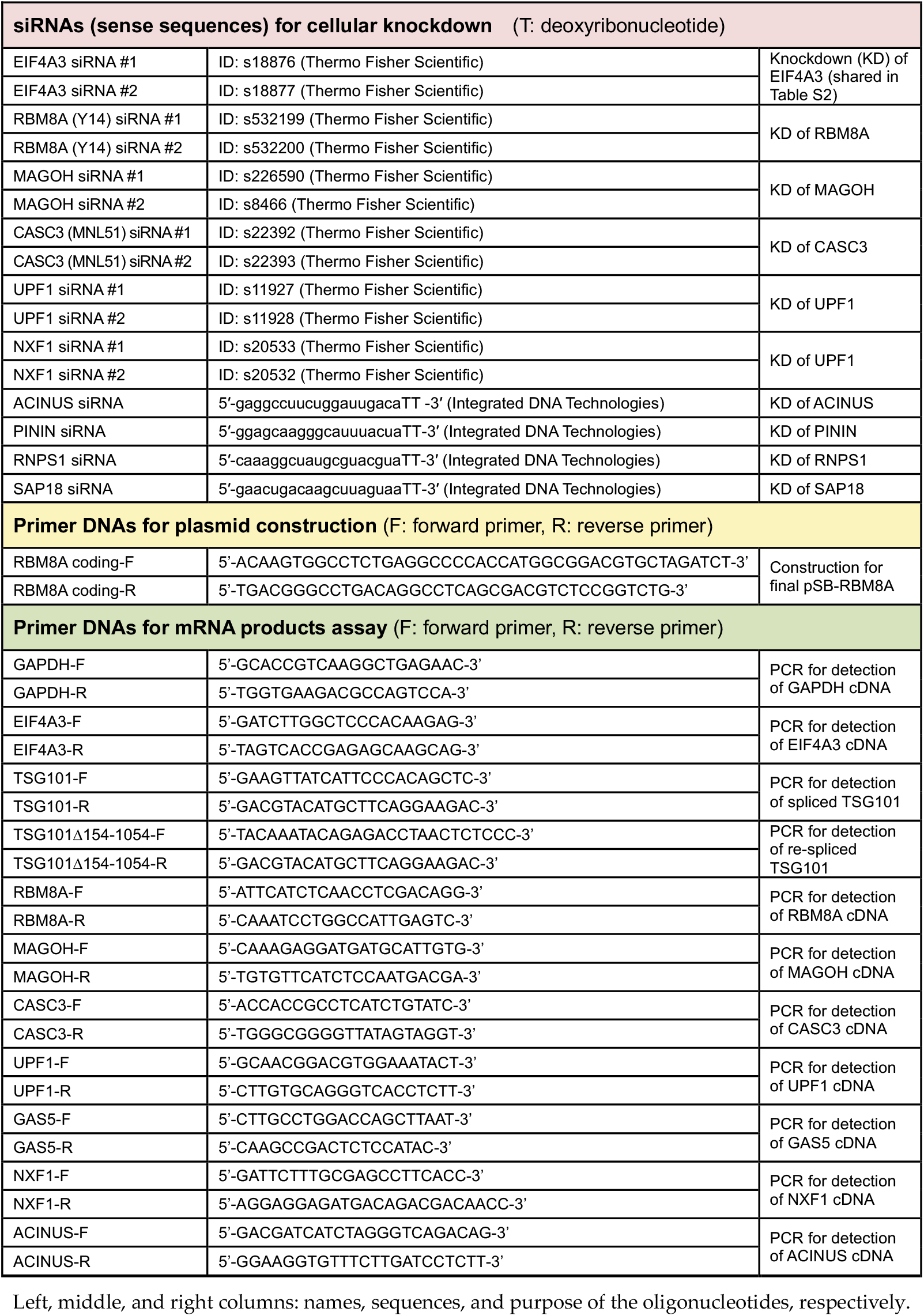
List of the synthetic oligonucleotides used in the experiments (see also Table S2).

**Figure S1.**
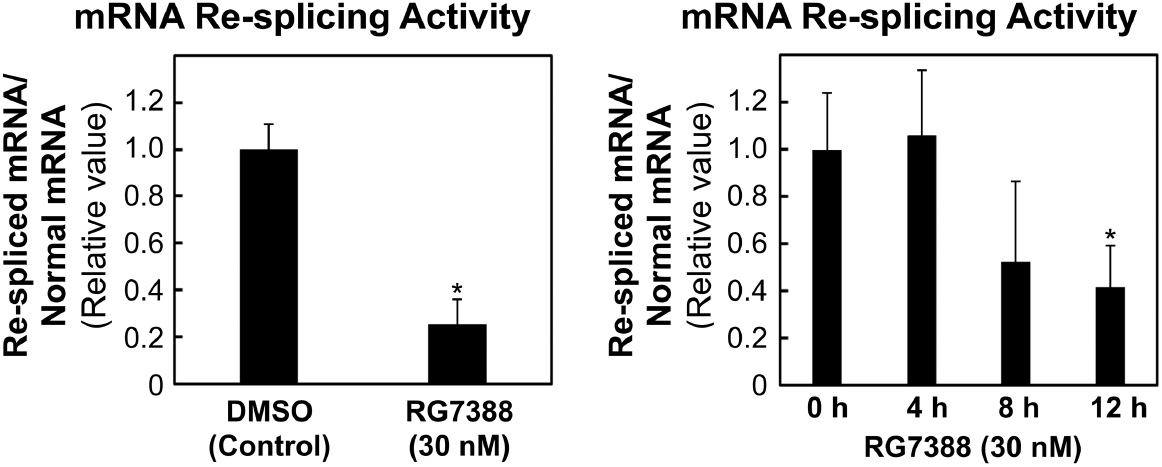
Activation of TP53 by the MDM2 inhibitor RG7388 induces the repression of TSG101 mRNA re-splicing activity: To evaluate the re-splicing activity with TSG101 mRNA, the extracted total RNAs from MCF-7 cells were analyzed by RT–qPCR using specific primer sets (see Figure 1a). The RG7388 concentrations and culture time after RG7388 addition were indicated. The histograms represent the means ± standard deviations of three replicates (\**p* < 0.05).

**Figure S2.**
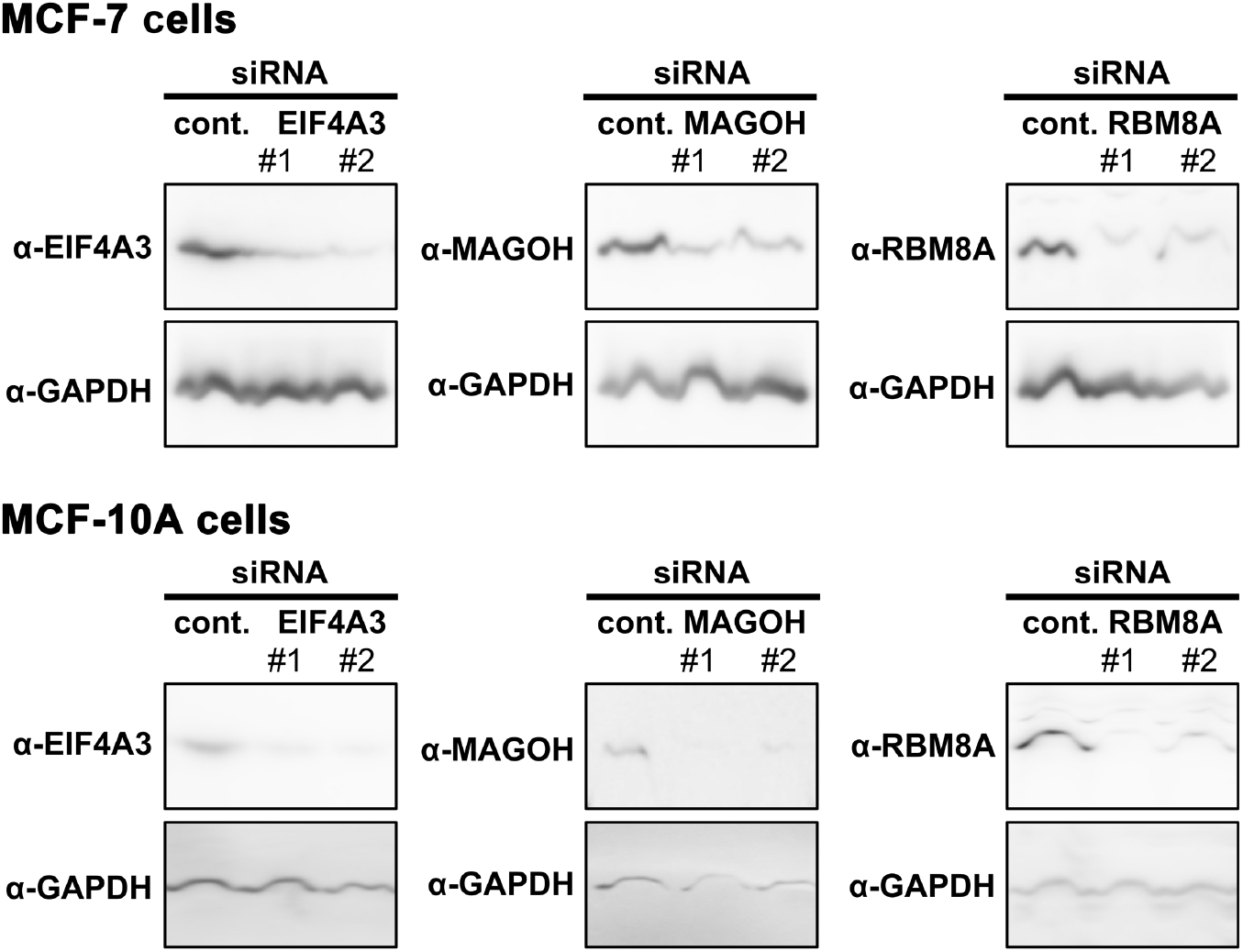
The endogenous EJC-core proteins are effectively depleted by the siRNA-mediated knock-down: MCF-7 and MCF-10A cells were transfected with siRNA targeting EIF4A3, RBM8A, MAGOH or control siRNA, and whole cell extracts were analyzed by immunoblotting with the indicated specific antibodies.

**Figure S3.**
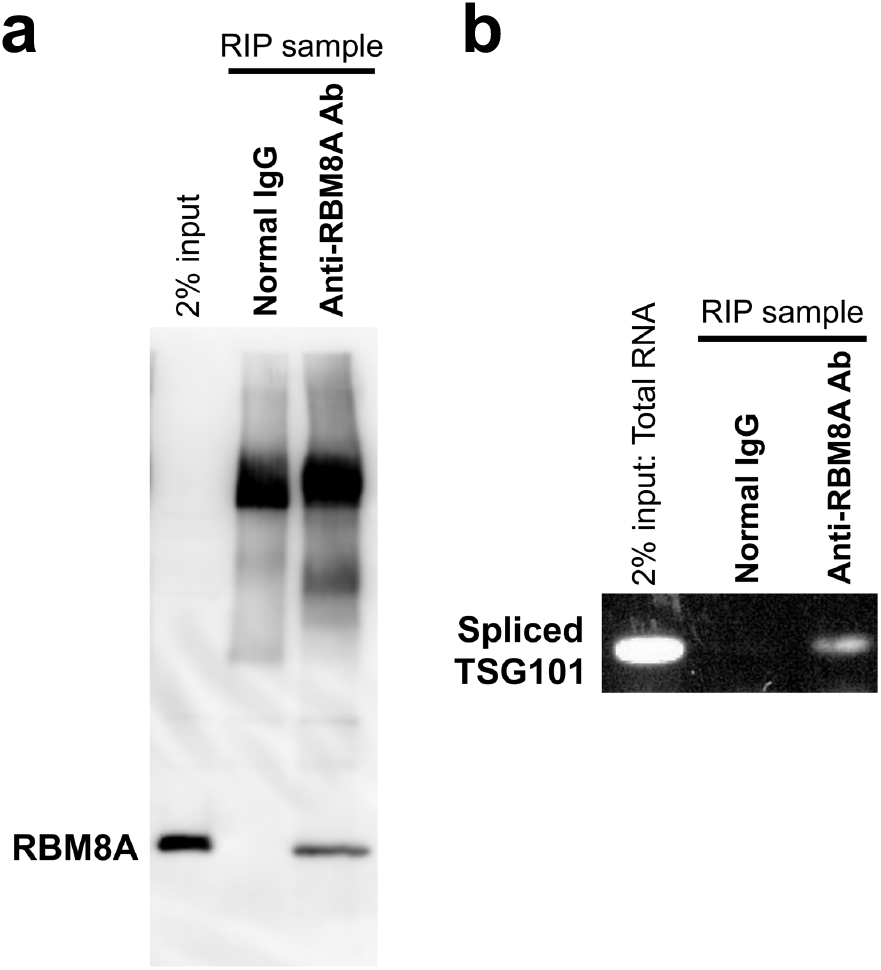
The EJC core factor RBM8A binds TSG101 mRNAs in MCF-7 cells: (**a**) RBM8A-bound mRNAs were separated from cell lysates of MCF-7 cells by RNA immunoprecipitation (RIP). Normal rabbit IgG was used as a negative control of the immunoprecipitation. Two independent RIP assays were performed and a representative immunoblot is shown here. (**b**) The RIP samples were analyzed by RT–PCR using a primer set targeting spliced TSG101 mRNA. The PCR products were visualized by agarose gel electrophoresis.

**Figure S4.**
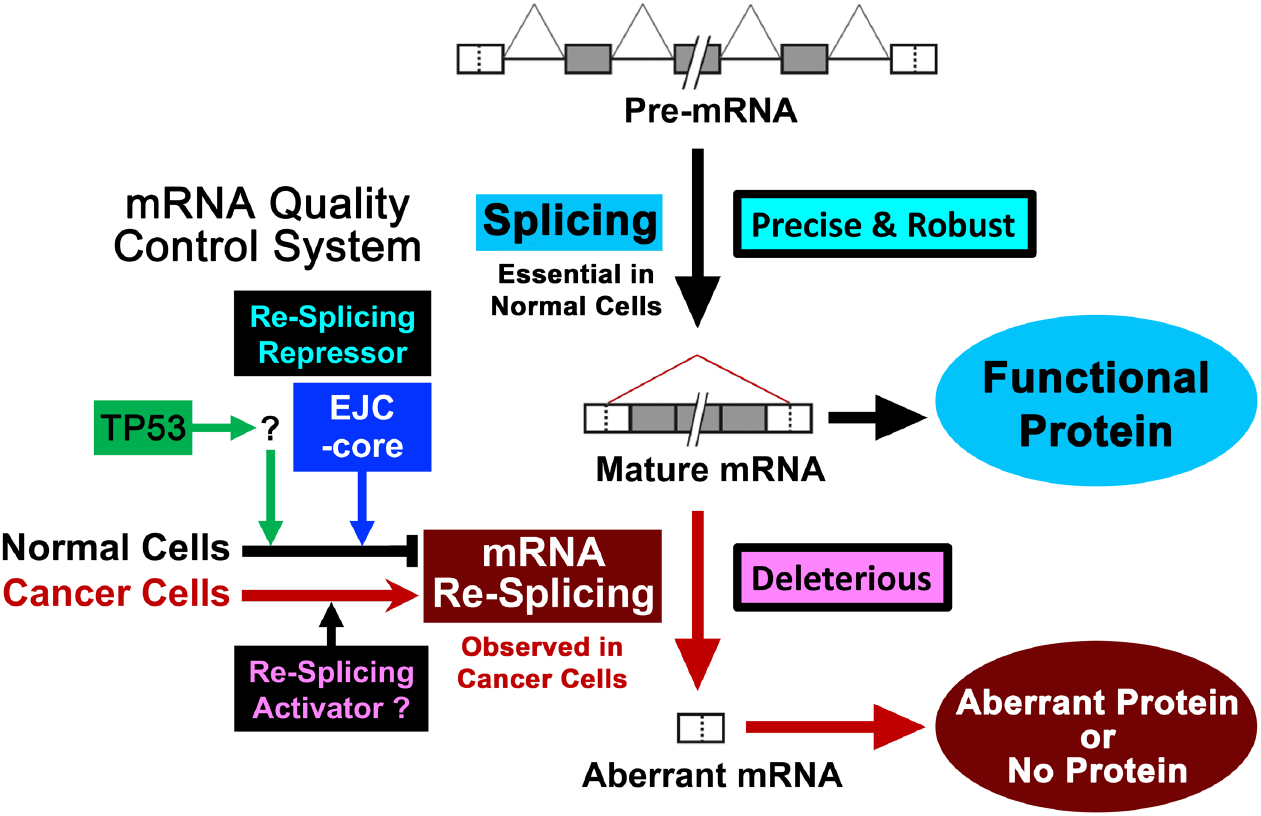
Model of the regulation system of cancer-specific mRNA re-splicing: TP53 represses mRNA re-splicing [10,15; see reference list in the main text], however, the downstream repressor has not been identified. Our discovery of EJC-core as a repressor of mRNA re-splicing implies a mechanism for terminating pre-mRNA splicing in normal cells, which is essential to maintain quality of mRNA.

**Figure S5.**
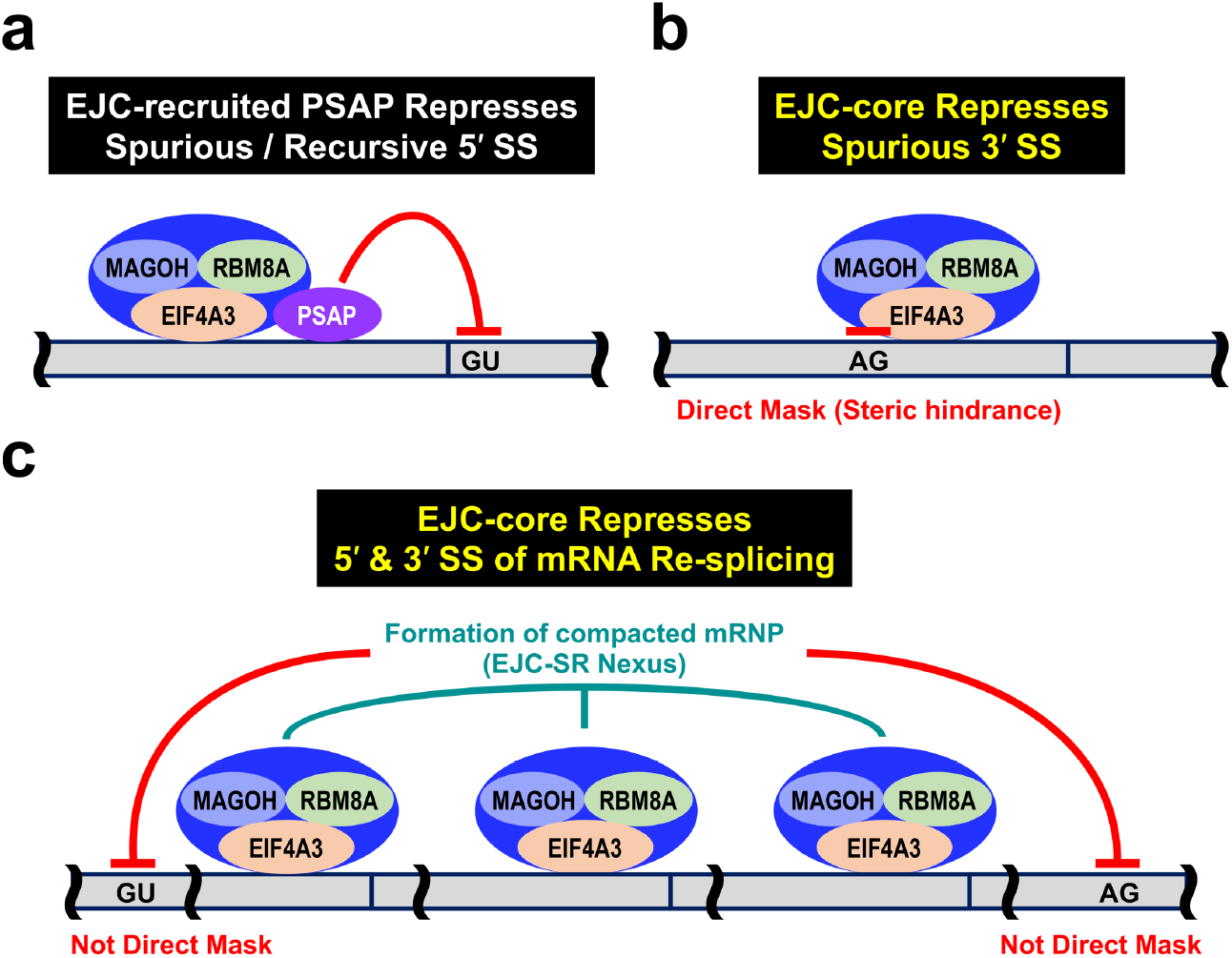
Similarity and distinction with other findings of the EJC-induced splicing repression: Recently, it has been reported that EJC deposition on mRNA represses nearby spurious splice sites [22; see reference list in the main text] and downstream recursive 5’ splice site [23]. (**a**) EJC-recruited PSAP complex (PNN, SAP18 and RNPS1) represses spurious 5’ splice site (SS) [22] and downstream recursive 5’ SS (but the requirement of PSAP is dependent on the substrate of recursive splicing) [23]. (**b**) On the other hand, nearby spurious 3’ SS is repressed by direct binding of the EJC-core, which causes steric hindrance [22]. (**c**) Here we showed that EJC-core represses exonic 5’ SS and 3’ SS used in mRNA re-splicing. Previous CLIP-Seq data suggest that the repression of mRNA re-splicing is not due to the direct binding of EJC-core on the 5’ SS and 3’ SS [24]. We propose that EJC-core deposition triggers the formation of compacted mRNP leading to the prevention of further re-splicing on mRNA (see Figure 6).

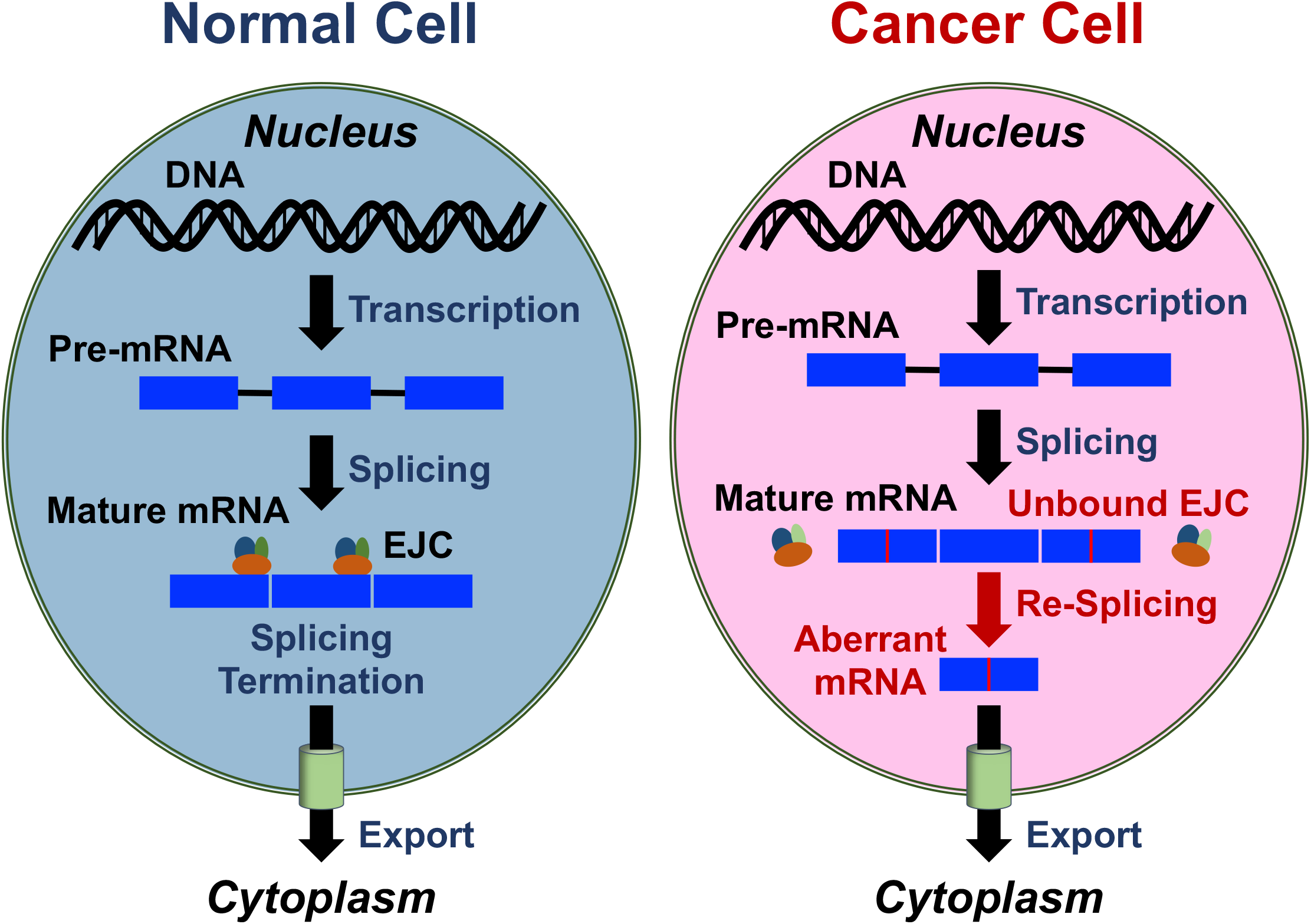

